# Gene tree patterns help answer: vicariance or dispersal?

**DOI:** 10.64898/2026.09.17.752061

**Authors:** Anna A. Nagel, Michael J. Landis, Fábio K. Mendes

## Abstract

Historical biogeography seeks to understand the drivers of species distributions over space and time. One question of interest is how, out of many possible ways, does geography drive speciation. Vicariance, where geographic barriers arise splitting populations and limiting gene flow, can lead to allopatric speciation. Founder events, where a small number of individuals disperse over a barrier, can similarly lead to allopatric speciation if the individuals remain isolated. Both of these scenarios can lead to identical ranges of and relationships between modern species. Classic biogeographic approaches often focus on the history of populations of one or few species on shallow time scales or multiple species on deep time scales. We argue that focusing exclusively on either end of this time spectrum misses a venue for investigating the biogeographic drivers of speciation, at least for certain speciation events. With simple coalescent simulations with multiple species, we show as a proof of concept that gene tree distributions vary predictably between vicariance and founder event speciation. The existence of predictable patterns warrants the development of new approaches that capitalize the gene trees to distinguish vicariance and dispersal as drivers of speciation.

---

Evolutionary studies often focus on either microevolutionary processes, operating within species at shallower timescales, or macroevolutionary processes, operating across multiple species at deeper timescales. Researchers have considered a wide variety of multiscale evolutionary phenomena, including quantitative trait evolution (Estes and Arnold, 2007; Mendes et al., 2018), molecular evolution (Muse and Gaut, 1994; De Maio et al., 2013; Ogilvie et al., 2022), and speciation (Fontaine et al., 2015; Meier et al., 2017; Wang and Hahn, 2018), to characterize how micro- and macroevolutionary phenomena jointly produce phylogenetic patterns (Rolland et al., 2023; Charlesworth et al., 1982). The geography of species, populations, and individuals also shapes how evolution produces phylogenetic patterns, as spatial context is highly relevant to adaptation (García-Ramos and Kirkpatrick, 1997), competition (Endler, 1977), demographic change (Hewitt, 2000), and speciation (Yamaguchi and Iwasa, 2013; Mayr, 1963). Taking this view, and building on pioneering work from the field of phylogeography (Avise, 2000; Knowles, 2009), we consider how gene tree topologies and coalescence times (Slatkin and Pollack, 2008; Hancock et al., 2022) obtained from species in a spatial radiation may contain useful information about geographical speciation. This information is, to our knowledge, rarely used in practice. In particular, different mechanisms for speciation, such as vicariance or founder events, can leave distinct signatures in gene trees that can aid inferences for how lineages originate in space. Our perspective piece first outlines how biogeography is typically considered from macro- and microevolutionary perspectives. Then, we show as a proof-of-concept that simple, biogeographically informed multispecies coalescent models lead to distinct gene trees patterns. Lastly, we suggest methodological developments that could capitalize on these patterns for inference and hypothesis testing.

## Phylogenetic Biogeography

Phylogenetic biogeography is chiefly concerned with how speciation, dispersal, and extinction are shaped by various geological, ecological, and climatological forces to produce phylogenetic trees and spatial patterns of species richness. A central question in phylogenetic biogeography concerns whether cladogenesis is more often the result of dispersal, in which a lineage disperses over an existing barrier and forms an isolated population that eventually becomes its own species, or the result of vicariance, whereby an emerging geographical barrier divides a widespread species into two new species. In contrast to the dispersalist view of diversification that prevailed into the 1960s (Matthew, 1915; Simpson, 1952; Hennig, 1966), vicariance biogeographers of the 1970s argued that one could test whether barrier-formation caused lineage divergence by simultaneously considering patterns of phylogenetic divergence, biogeographic disjunction, and earth history (Platnick, 1978; Rosen, 1978). In the same breath, some vicariance biogeographers also held that speciation-through-dispersal (e.g., through founder events (Mayr, 1954) or long-distance dispersal (Carlquist, 1966) could occur anywhere and anytime, making it unfalsifiable as a hypothetical mechanism for speciation (Platnick and Nelson, 1978; Croizat et al., 1974). Dispersalist explanations grew in favor since the 2000s, fueled by new molecular sequencing technologies and phylogenetic methods, with hundreds (if not thousands) of new studies showing that dispersal over barriers is necessary to explain diversification patterns (Crisp et al., 2011; Matzke, 2014; De Queiroz, 2014). In general, however, these findings generally derive from ancestral range estimates that are informed only by species-level phylogenetic relationships, divergence times, and biogeographic occurrences (Ree et al., 2005). While more complex analyses can make use of other data, such as the times of paleogeological dynamics (Ree and Smith, 2008), regional features that shape biogeographic rates (Landis et al., 2022), or fossil evidence (Wood et al., 2013), it remains difficult to interpret the underlying mode of speciation from ancestral state patterns alone. In addition, since classical approaches for phylogenetic biogeography assume each species possesses exactly one range (a set of occupied regions), they have limited information to confidently reconstruct the mode of speciation for any particular divergence event.

### Phylogeography

Phylogeography, as originally defined, is the study of gene tree patterns in a spatially explicit context (Avise et al., 1987; Avise, 2000). Typically phylogeographic studies focus on either populations within the same species or from a small number of species, often pairs of sister taxa. In the former case, single-species studies focus on inferring changing demographic histories through time to differentiate among alternative hypotheses often informed by geological histories, paleoclimate data, pollen cores, ancient DNA, or fossils (Hickerson and Meyer, 2008; Gavin et al., 2014; Ortego and Knowles, 2022). This may include questions such as how population connectivity and population sizes changed over the glacial cycles to modern day, or whether sea level changes can account for the current population structure within a species (Pelletier and Carstens, 2014; Dantas-Queiroz et al., 2023). In the latter case, studies across species often focus on testing whether a particular geologic event caused multiple, often distantly related species to begin diverging at the same time (Oaks et al., 2013, 2022; Busschau et al., 2022). However, caution must be taken to avoid equating gene tree divergence times with species tree divergence times, as this can bias estimates and undermine hypothesis testing (Edwards and Beerli, 2000). For example, changing ocean levels are implicated in splitting islands and the populations on them, creating distinctive patterns of genetic variation, and leading to vicariant speciation (Knowles and Maddison, 2002). But it could also be that dispersal to islands and subsequent speciation after sea level rise or speciation prior to sea level rise could give rise to the same species distributions. Discriminating between very simple alternative hypotheses, such as these, can be difficult due to the complex relationship between gene trees and population histories, particularly since the gene trees themselves are estimates. The challenges grow even greater when a variety of geologic factors are thought to be important in a region (e.g., Brown et al., 2013).

### Coalescent Theory

Many advances in phylogeography were propelled by developments in coalescent theory. The coalescent model describes the distribution of gene trees underlying a set of samples from a population (Kingman, 1982a,b). In a simple case, consider a diploid population of *N* individuals with non-overlapping generations (i.e., a Wright-Fisher population; Fisher, 1930; Wright, 1931). Traveling backwards in time, two genes will eventually meet in their most recent common ancestor, an occasion known as a coalescence event. The waiting time until the coalescence event reduces the number of lineages from *n* to (*n −* 1) samples is

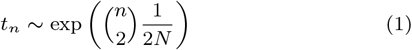

Through this formulation, the distribution of coalescence times allows the effective population size (scaled by the mutation rate) to be estimated from genetic data.

Coalescent theory has been extended to allow for varying population size through time (Slatkin and Hudson, 1991; Griffiths and Tavaré, 1994; Li and Durbin, 2011; Schiffels and Durbin, 2014; Pybus et al., 2000; Strimmer and Pybus, 2001), and explicitly account for current and ancestral population structure (Nielsen and Wakeley, 2001; Hey and Nielsen, 2007). The multispecies coalescent (MSC) models how species divergence shapes the coalescent process (Takahata, 1989; Rannala and Yang, 2003). Under the MSC, the genetic samples within each population (species) evolve independently according to the Kingman coalescent until a divergence time is reached (backward in time). Then, all surviving lineages from the descendant populations become lineages in the parent population. Inference methods have also been developed for coalescent models with gene flow (e.g., through the process termed migration by population geneticists; Beerli and Felsenstein, 1999, 2001), many of which have been incorporated into MSC models (Flouri et al., 2020, 2023). Coalescent models have also been applied in a spatial context to consider how geographically structured populations shape gene trees (Slatkin and Pollack, 2008; Obbard et al., 2012; Hancock et al., 2022). These extensions allow for simulation of and inference about a wide variety of biogeographical questions on demography, population continuity, structure, and divergence.

### Limitations of Coalescent Theory

A key limitation when using coalescent-based methods to estimate population size (*N* in Eqn. 1) is that the signal from data for such inferences diminishes at deeper timescales. Adding more samples (increasing *n* in Eqn. 1) increases the rate coalescent of events, so the additional samples tend to coalesce quickly, and thus generate little additional signal about more-ancient events. More generally, the expected coalescence time for the MRCA of an entire population is 4*N*, with all coalescent events associated with descendants of the MRCA being more recent. This is to say that, under certain circumstances, there are fundamental limits to how much can be learned from population genetic data alone.

However, this limitation for estimating population sizes at deeper times only applies to samples from a single population, but not to samples from two or more divergent populations under the MSC. Gene lineages must belong to the same population to coalesce, and lineages from different populations may eventually coalesce in two ways. First, gene flow (e.g., via migration) enables gene lineages to coalesce between divergent populations, whereby one lineage enters the alternate population. Second, gene lineages from divergent populations that do not experience gene flow may coalesce only ‘after’ (going backwards in time) the ancestral population exists. It is only possible to infer demographic histories in terms of *N* during time periods when there are coalescent events, and population divergence events delay coalescence until the divergence time (again, backwards in time). Under the MSC, demographic information for an ancestral population can be inferred at timescales much older than is possible from one population alone. That said, common usage of the MSC assumes population sizes for each ancestral species is constant over time. In principle, more complex demographic histories pertaining to cladogenesis, such as population bottlenecks and/or subdivisions concomitant with speciation, could be inferred for ancestral populations.

## Geographical speciation shapes gene tree distributions

Common verbal models of allopatric speciation lead to distinct predictions about demographies and population structure through time. Consider a typical vicariance scenario from a population genetic perspective (Slatkin and Pollack, 2008). Initially a species occupies a large area, i.e., it is widespread. A physical barrier to dispersal (and thus to gene flow) arises, such as rising sea levels splitting islands, dividing the population into two. There may be some gene flow as the barrier emerges, even as genetic differences accumulate in each population. Eventually, the populations become either sufficiently diverged or the barrier sufficiently strong such that gene flow ceases, leading to speciation. This pattern may be repeated again in the daughter species (Fig. 1a). In contrast, in founder event speciation, a very small number of individuals disperses from a larger population over a barrier, such as across an ocean to an island, and then experience substantial population growth (Obbard et al., 2012). There may be limited gene flow with the ancestral population for some period of time before speciation completes. This may be repeated across multiple barriers or areas, such as across a chain of islands (Fig. 1b). Both of these scenarios may leave identical modern ranges of species (Fig. 1a,b) and species trees (Fig. 1c).

**Figure 1.**
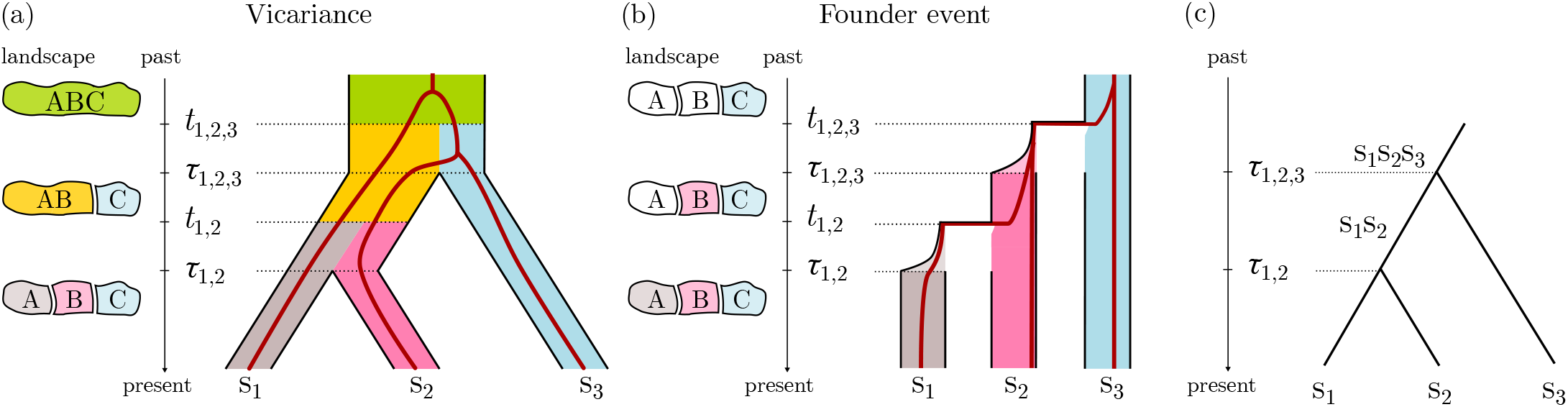
(a) Initially a diploid population of size 3*N* distributed across a large area (green). A barrier begins to form at time *t*_1,2,3_, separating the landscape into regions AB and C and creating two populations of size *N*_1,2_ (yellow) and *N*_3_ (blue), where *N*_1,2_ + *N*_3_ = 3*N*. Migration between populations occurs with a symmetric rate of *m*_*v*_, where the rate refers to the proportion of the population replaced each generation in forward time. At time *τ*_1,2,3_ gene flow ceases. A second barrier arises at time *t*_1,2_ that splits region AB into A and B, leading to two populations of size *N*_1_ (gray) and *N*_2_ (pink) where *N*_1_ + *N*_2_ = *N*_1,2_ with symmetric gene flow between them at rate *m*_*v*_. At time *τ*_1,2_, gene flow ceases. (b) Initially a single diploid population of size *N* is in region C. At time *t*_1,2,3_, *N*_*b*_ individuals leave the population and colonize region B. The population grows exponentially, reaching size *N* at time *τ*_1,2,3_. The parent population grows at the same rate until it reaches its original size, *N*. Symmetric migration occurs at rate *m*_*f*_ until speciation completes at time *τ*_1,2,3_. At time *t*_1,2_, *N*_*a*_ individuals leave the population s_2_ and colonize region A. The new population grows exponentially and reaches size *N* at time *τ*_1,2_. As before, the parent population grows at the same rate until it reaches its original size, *N*. Migration occurs at *m*_*f*_ until *τ*_1,2_. (c) Both panels (a) and (b) give rise to the same species tree with three species, s_1_, s_2_, and s_3_, and the same divergence times, *τ*_1,2_ and *τ*_1,2,3_. The common ancestor s_1_ and s_2_ is called s_1_s_2_. The population ancestral to all species is labeled s_1_s_2_s_3_.

If these biogeographic scenarios produce distinct distributions of gene trees, they could be useful for inferring the underlying mode of speciation (Knowles and Maddison, 2002). To illustrate this point, we formalize the above descriptions for vicariance and founder event speciation within the framework of the MSC. We consider four scenarios along a parameter gradient going from “Realistic vicariance” (scenario 1) to “Realistic founder event” (scenario 4), with “Founder event-like vicariance” (scenario 2) and “Vicariance-like founder event” (scenario 3) in between. In scenario 2, a physical barrier is assumed to split a widespread population into two sub-populations of very unequal size, while scenario 3 consists of a mild bottleneck during a founder event with migration rates higher than scenario 4’s. For each of the four scenarios, we simulate 100 independent coalescent histories (i.e., gene trees), with one gene sample per locus for each of three species labeled ‘s_1_’, ‘s_2_’ and ‘s_3_’ (Fig. 1). Exact parameter values used in the scenarios above and other scenarios we investigated can be found in the supplement.

The gene trees from the “Vicariance” and “Founder event” scenarios differ substantially, both in coalescence times and topologies. The “Vicariance” scenario has the most discordant gene trees (the gene tree does not topologically match the species tree due to incomplete lineage sorting) and older coalescence times on average (Fig. 2a) (Slatkin and Pollack, 2008). In contrast, the “Founder event” scenario has little gene tree discordance and much younger coalescent times on average, since the bottleneck almost guarantees the coalescent time of samples from s_1_ and s_2_ is younger than the founder event (Obbard et al., 2012). Since the divergence times are shared by all scenarios, their distributions of the minimum coalescence times are similar, but the variance of the coalescence times and the maximum coalescent times are much higher in the “Vicariance” than in the “Founder event” scenario (Fig. 2b). This is because in the “Founder event” scenario, coalescent times are clustered around the bottleneck. In between these two scenarios is “Vicariance-like founder event”, where the weaker bottleneck causes only some lineages to coalesce before reaching the root population. Lastly, the “Founder event-like vicariance” scenario has similar levels of discordance and average node as the “Vicariance” scenario, due to similarly large population size of s_1_s_2_. However, the minimum coalescence times and maximum coalescent times tend to be somewhat larger, due to the population size of s_1_s_2_ being somewhat larger than in the “Vicariance” scenario.

**Figure 2.**
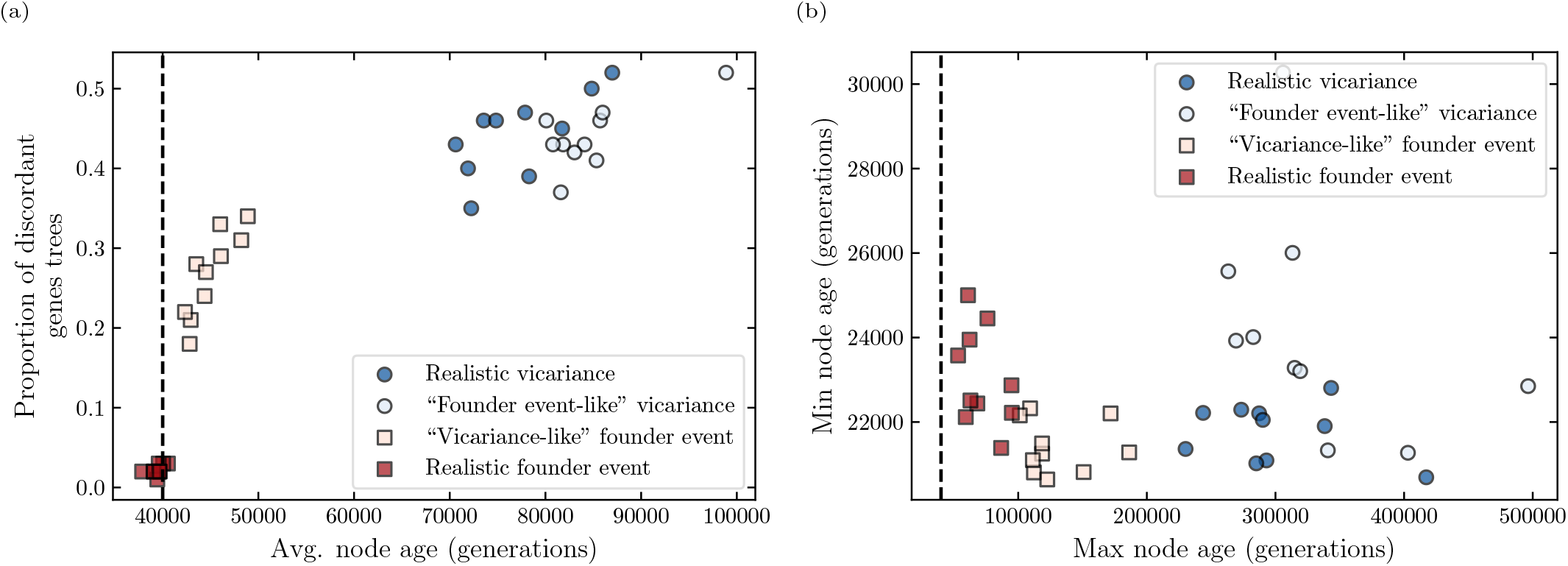
Distributions of coalescent times and discordant gene trees. (a) The x-axis shows the average coalescent time of one sample from each of s_1_ and s_2_. The y-axis shows the proportion of discordant gene trees (topology does not match the species tree topology). (b) The minimum coalescence time by the maximum coalescence time of the samples from s_1_ and s_2_ for each simulation. Gene trees were simulated with msprime (Baumdicker et al., 2022). Each simulation has one sample from each species and 100 gene trees. There are 10 replicates of each scenario. The dashed line shows *τ*_1,2,3_ = 4 × 10^4^, which is the time the population decrease (going backward in time) begins in the founder event and migration begins between subpopulations. The other parameters are *τ*_1,2_ = 2 × 10^4^, *t*_1,2_ = 3 × 10^4^, *t*_1,2,3_ = 5 × 10^4^. For the realistic vicariance scenario, *N* = *N*_1_ = *N*_2_ = *N*_3_ = 10^4^, and *m*_*v*_ = 0.0001. For the “founder event-like” vicariance, *N*_1_ = 24, 300, *N*_2_ = 2, 700, *N*_3_ = 3, 000 and *m*_*v*_ = 5 × 10^*−*6^. For the realistic founder scenario, *N* = 10^4^, *b* = 0.001, and *m*_*f*_ = 5 × 10^*−*6^. For the “vicariance-like” founder event, *N* = 10^4^, *b* = 0.33, and *m*_*f*_ = 0.0001.

The results discussed above only touch on the older of the divergence events (Fig. 1). However, gene trees can contain information about the younger of the two divergence events as well (Supplement 3.4, Fig. S7,8). While our simulations are not exhaustive, the general patterns reported above seem to be relatively insensitive to specific parameter choices as long as the scenarios retain some similarities to ensure they are comparable (e.g., the same divergence times across all scenarios; Supplement 2 and 3, Figs. S2-S8).

### Empirical Considerations

Even if different scenarios for geographical speciation can produce distinct gene tree distributions, should we expect that similar patterns will be apparent under broader range of empirical conditions? For many highly studied radiations, species have population sizes and relative divergence times such that their genetic data are likely to contain demographic and biogeographic signatures from the time of speciation (Table 1). In particular, when the branch lengths in generations are the same order of magnitude as the population size, there are discernible differences in gene tree distributions between the four demographic models (Fig. S2). While table 1 specifically focuses on rapid and recent radiations, gene tree patterns could be expected to also apply to older radiations (e.g., Fig. S3) or individual branches within a phylogeny that are short relative to the population size.

**Table 1.** Population genetic parameters for several radiations. When the population size for the species is similar to the branch lengths measured in generations, there will likely be information in genetic data regarding speciation histories for at least some branches in the phylogeny. Branch lengths in generations were found by dividing the branch lengths in years by the generation time. The very small population size for one of Darwin’s Finches reflects a recent population collapse. For the African Cichlids average branch lengths in years were found by dividing the range of the age estimates of the radiation by the number of branches. The range reflects the uncertainty in root age estimates, not the actual branch-length range in the tree. The population sizes for the Hawaiian honeycreepers are historical sizes, not recent census sizes. The branch lengths are from a phylogeny of 19 extant and recently extinct species, not the entire radiation of more than 50 species. Very small modern census sizes is not generally expected to negatively impact the ability to infer population sizes for ancestral populations.

| Clade | number of species | population size (per species) | branch lengths (years) | generation time | approximate branch lengths (generations) | citations |
| --- | --- | --- | --- | --- | --- | --- |
| Darwin's Finches | 14 or 17 | 80–600,000 | <10,000-100,000 within main radiation | ~5 years | <2,000-20,000 | Lamichhaney et al. (2015) reviewed in Hedrick (2019) |
| African Cichlids |  | 1,000-100,000 |  | ~3 years |  | Malinsky et al. (2015) |
| Lake Tanganyika | ~250 |  | 20,000-60,000 |  | 7,000-20,000 | Marques et al. (2025) |
| Lake Malawi | ~900 |  | 550-3,000 |  | 200-1,000 | Loh et al. (2013) |
| Lake Victoria | ~600 |  | 130-830 |  | 40-280 | and citations therein |
| Hawaiian Honeycreepers | 17 extant | 40,000-150,000 | 0.2-2.5 million | ~4 years | 50,000-625,000 | Lerner et al. (2011) Kyriazis et al. (2025) |

## Unifying biogeography across scales

The idiosyncratic approaches to biogeography in different fields can make it difficult to translate across temporal, spatial, and taxonomic scales. Moreover, thinking only within species or across species with species trees disregards the signatures that population genetic processes leave deep within the gene trees. As suggested by our initial investigation, distinct biogeographic hypotheses such as vicariance and founder-event speciation predict visibly different gene tree distributions. Such differences should not depend on absolute divergence times (Oliver, 2013) – that is, on whether speciation events are recent or ancient – but on their temporal spacing. We propose that studying distributions of gene trees in the context of speciation will help biogeographers disentangle competing hypotheses regarding modes of speciation, potentially for ancient divergence events, in cases that can be difficult for existing methods.

Fitting multispecies coalescent models of biogeography to empirical datasets will require the development of new methods. To adequately capture both the complexity of the models and uncertainty, ideally such approaches would be built on probabilistic frameworks that jointly model gene flow and demographic parameters across species within a full Bayesian setting (Flouri et al., 2023; Wen et al., 2016). The models of vicariance and founder events could in principle be distinguished using Bayes factors or model adequacy. Other factors, such as the temporal placement of such events, together with the appearance of barriers, could be a function of paleogeography (Lichter-Marck et al., 2025), as certain areas may become inhabitable or more likely to receive migrants at particular times.

In practice, however, fully Bayesian approaches will be computationally prohibitive for any but the smallest phylogenies. Alternative methods rely heavily on simulation. Approximate Bayesian computation (ABC), which has a well-developed track record in phylogeography (Hickerson et al., 2006; Cornuet et al., 2008; Pelletier and Carstens, 2014) and could be naturally extended to gene-tree distributions as the relevant summaries. Machine learning approaches offer a complementary path, with convolutional neural networks having recently shown promise for phylogeographic model selection (Fonseca et al., 2021) and broader population genetic inference (Korfmann et al., 2023). Many radiations of interest in historical biogeography are much older than the examples discussed here, with branch lengths much larger than the effective population sizes. In this case, using machine learning or ABC may return noise rather than true signals of biogeographic history. Rather than let this prevent method development, rough estimates of branch lengths and populations sizes should suggest whether gene tree based approaches are expected to give true signatures of the data.

Several factors will likely complicate empirical analyses. Most importantly, the branch lengths relative to the population size determine whether different speciation scenarios are expected to lead to different patterns in the gene trees. This approach will likely work better when the branches are short or the population sizes are large, such as in radiations. Our results hinge on having predictable demographic patterns for different drivers of speciation. Population sizes can nonetheless vary dramatically overtime within a population, independent of speciation, which can shift the distribution of gene trees to such a degree that they may no longer be informative for the purposes discussed here. Additionally, population structure and demography can be confounded, at least in some cases (Chikhi et al., 2010; Heller et al., 2013). Whether this will obscure patterns related to speciation histories remains to be determined.

## Data availability

Simulation scripts are available at https://github.com/mendeslab/geogenetree/tree/extensions.

