## supplemental material for "Gene tree patterns help answer: vicariance or dispersal?"

Supplement

Anna A. Nagel, Michael J. Landis, and Fábio K. Mendes

#### 1 Motivation

The simulations discussed in the main text suggest that gene distributions can help identify vicariance versus dispersal events leading to speciation. However, the observed gene tree patterns could be due to idiosyncrasies of the model or parameters of choice, which would limit the applicability of the gene tree-based strategy we propose for studying the biogeography of speciation. We thus conducted additional simulations to examine whether the gene-tree distributions generated under distinct biogeographic hypotheses remained distinguishable under a wide range of parameter values and modeling choices. Should that be the case, it suggests future methods in this vein – that is, which leverage the distribution and characteristics of gene trees – may be broadly applicable. Here, the goal was to retain key aspects of the scenarios investigated in the main text, while relaxing some assumptions.

#### 2 Extended Model

We generalize the coalescent model from the main text (Fig. 1) to allow for fewer symmetries in the parameter values. Then, this model is used to simulate gene trees. As usual, coalescent models are in backward time, with time measured *in generations* and flowing from the present (time zero) towards the past. Note that in what follows, however, we describe the model events in forward time unless specified otherwise.

In all the scenarios we examine, the species tree has the same topology, $((\mathrm{s}_1,\mathrm{s}_2),\mathrm{s}_3))$, with terminal nodes (extant species) $\mathrm{s}_1$, $\mathrm{s}_2$, and $\mathrm{s}_3$. The species that is ancestral to $\mathrm{s}_1$ and $\mathrm{s}_2$ is referred to as $\mathrm{s}_1\mathrm{s}_2$. This species exists between times $\tau_{1,2,3}$ (when this species splits from its sister) and $\tau_{1,2}$ when it speciates, giving rise to species $\mathrm{s}_1$ and $\mathrm{s}_2$. There is population substructure within $\mathrm{s}_1\mathrm{s}_2$ between times $t_{1,2}$ and $\tau_{1,2}$. Similarly, the root species is referred to as $\mathrm{s}_1\mathrm{s}_2\mathrm{s}_3$, existing from some untracked time in the past until $\mathrm{t}_{1,2,3}$ (the root node's age). The root species has population substructure between $t_{1,2,3}$ and $\tau_{1,2,3}$. Note that the durations of species $\mathrm{s}_1\mathrm{s}_2$ and $\mathrm{s}_1\mathrm{s}_2\mathrm{s}_3$ are not affected by their populations being structured. In all models described below, all species are diploid and migration rates $m_{i \to j}$ denote the proportion of species $j$'s population that is replaced by species $i$'s population in each generation in forward time.

Because the models described below are standard population genetics coalescent models, there is no explicit notation for referencing biogeography. Biogeographic ranges and events are implied by changing population structure and connectivity (i.e., migration), amounting to the same biogeographic scenarios described in the main text. In both the vicariance and founder event scenarios, all three extant species occur in only one disjoint region each. To match the main text's vicariance event model, each ancestral population would occupy multiple locations. In turn, the founder event model has each ancestral population occupy a single region.

**Vicariance**. In forward time, a starting species $\mathrm{s}_1\mathrm{s}_2\mathrm{s}_3$ has a panmictic population of size $3N$ ($N_{1,2,3} = 3N$; Fig. S1a). This panmictic population is subdivided into two populations at time $t_{1,2,3}$, when they begin to diverge from one another; these populations are $\mathrm{s}_1\mathrm{s}_2$ of size $N_{1,2}$ and $\mathrm{s}_3$ of size $N_3$, where $N_{1,2} + N_3 = 3N$. This event represents the effect of a *geographic barrier* appearing at time $t_{1,2,3}$, which *limits gene flow* between the two populations and results in the subdivision. After the subdivision, migration is assumed to occur at rate $m_{1,2 \to 3}$ from population $\mathrm{s}_1\mathrm{s}_2$ to population $\mathrm{s}_3$ forward in time. Similarly, migration from population $\mathrm{s}_3$ to population $\mathrm{s}_1\mathrm{s}_2$ occurs at rate $m_{3 \to 1,2}$. At time $\tau_{1,2,3}$ gene flow ceases between $\mathrm{s}_1\mathrm{s}_2$ and $\mathrm{s}_3$, at which point speciation is assumed to be complete (i.e., in the species tree, $\tau_{1,2,3}$ can be seen as a divergence or speciation time, or as the age of the root node).

At time $t_{1,2}$ a similar series of events take place. Population $\mathrm{s}_1\mathrm{s}_2$ is subdivided, creating populations $\mathrm{s}_2$ and $\mathrm{s}_1$ of size $N_1$ and $N_2$, respectively, where $N_1 + N_2 = N_{1,2}$. Migration occurs at rate $m_{1 \to 2}$ from population $\mathrm{s}_1$ to population $\mathrm{s}_2$, and from population $\mathrm{s}_2$ to population $\mathrm{s}_1$ at rate $m_{2 \to 1}$. Lastly, at time $\tau_{1,2}$ gene flow ceases and these two populations are assumed to be different species.

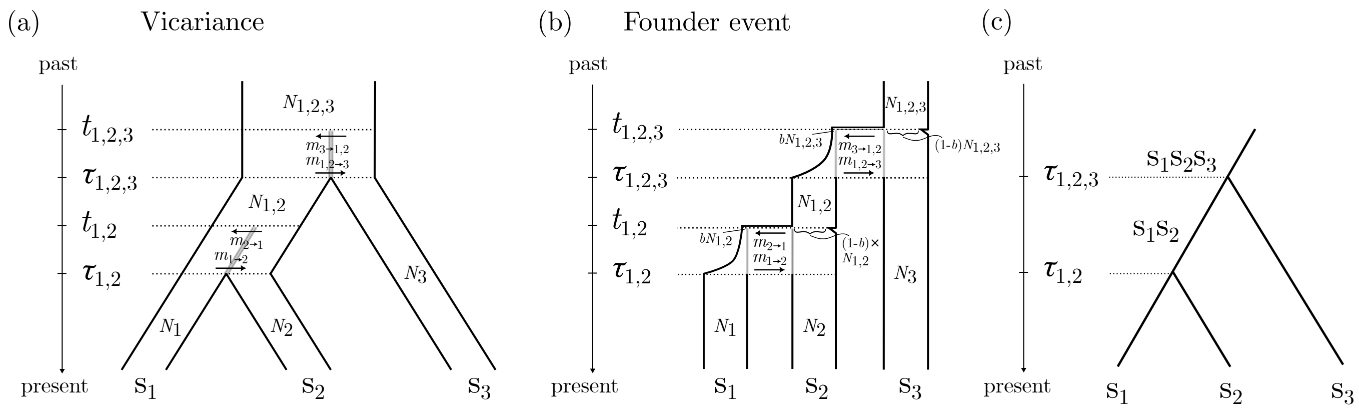


Figure S1: Population structuring times are denoted by $t$, where is speciation completion times are denoted by $\tau$. Population sizes are denoted by $N$, migration rates by $m$, and bottleneck intensity by $b$. See text for further details. (a) Vicariance scenario drawn as a MSC tree with migration. (b) Founder event scenario drawn as MSC tree with migration. (c) Species tree represented in both (a) and (b), but drawn as a simple chronogram.

**Founder event.** In forward time, a starting species $s_1s_2s_3$ has a panmictic population of size $N_{1,2,3}$ (Fig. S1b). Upon a founder event at time $t_{1,2,3}$, $bN_3$ individuals migrate out of population $s_1s_2s_3$ and found a new population, $s_1s_2$. The remainder of population $s_1s_2s_3$ is now called $s_3$. Population $s_1s_2$ grows exponentially, reaching size $N_{1,2}$ at time $\tau_{1,2,3}$. Meanwhile starting from time $t_{1,2,3}$, population $s_3$ also grows exponentially (at the same rate) until it reaches $N_3$, which equal $N_{1,2,3}$, the size it had prior to the foundation of population $s_1s_2$. Migration occurs at rate $m_{1,2\to3}$ from population $s_1s_2$ to population $s_3$ forward in time and at rate $m_{3\to1,2}$ from population $s_3$ to population $s_1s_2$. At time $\tau_{1,2,3}$, gene flow between populations $s_1s_2$ and $s_3$ stops completely and these two populations are considered different species (i.e., in the species tree, $\tau_{1,2,3}$ can be seen as a divergence or speciation time, or as the age of the root node). Note that from time $t_{1,2,3}$ to time $\tau_{1,2,3}$, species $s_1s_2s_3$ is considered widespread *across a barrier to dispersal that limits gene flow* to some degree.

Upon a second founder event at time $t_{1,2}$, a similar series of events take place. $bN_{1,2}$ individuals leave population $s_1s_2$ and found population $s_1$. The remainder of population $s_1s_2$ is now called $s_2$. Population $s_1$ grows exponentially and reaches size $N_1$ at time $\tau_{1,2}$. As before, the other population, $s_2$, grows at the same rate until it reaches size $N_2$, which equal $N_{1,2}$, the size it had prior to the foundation of population $s_1$. Migration occurs at rate $m_{1\to2}$ from population $s_1$ to population $s_2$, and at rate $m_{2\to1}$ in the opposite direction. Gene flow ceases at time $\tau_{1,2}$, at which point these two populations are considered two different species.

### 3 Extended Simulations Results

Simulations were conducted with MSPRIME (Baumdicker et al., 2022) to examine whether the distinct distributions of gene trees for each scenario were robust to the specific parameter and modeling choices. The goal was to retain the key aspects of each scenario (realistic vicariance, realistic founder, founder event-like vicariance, and vicariance-like founder), while relaxing some assumptions. In comparison to the main text, for example, the simulations consider different parameter settings (tables 1 to 5) changing:

- internal branch lengths (parameter settings 2-4),
- terminal branch lengths (parameter settings 5-7),
- population size of $s_1s_2$ (parameter settings 8), and
- migration rates (parameter settings 9-12).

For every simulation, 100 gene trees were drawn and each scenario replicated ten times (all the plots have 10 points for each scenario). Unless otherwise stated, one sample was drawn for each species. All parameters for the simulations are shown in tables 1 to 5. In some cases, parameter settings were repeated for one of the scenarios to compare to cases where the parameter settings were different in a different scenario. For example, parameter settings 1 and 8 are the same for the vicariance scenario but different for the founder event scenario. In the "vicariance-like founder event" scenario, the migration rates were the same as for the vicariance event scenario and the bottleneck

| parameter settings | figure | $N_1$ | $N_2$ | $N_3$ | $m_{1\to 2}$ | $m_{2\to 1}$ | $m_{1,2\to 3}$ | $m_{3\to 1,2}$ |
|---|---|---|---|---|---|---|---|---|
| default/1 | 1, S7 | $10^4$ | $10^4$ | $10^4$ | $10^{-4}$ | $10^{-4}$ | $10^{-4}$ | $10^{-4}$ |
| 2 | S2a,b | - | - | - | - | - | - | - |
| 3 | S2c,d | - | - | - | - | - | - | - |
| 4 | S2e,f | - | - | - | - | - | - | - |
| 5 | S3a,b | - | - | - | - | - | - | - |
| 6 | S3c,d, S8 | - | - | - | - | - | - | - |
| 7 | S3e,f | - | - | - | - | - | - | - |
| 8 | S4 | - | - | - | - | - | - | - |
| 9 | S5a,b | - | - | - | 0 | 0 | 0 | 0 |
| 10 | S5c,d | - | - | - | $10^{-3}$ | $10^{-3}$ | $10^{-3}$ | $10^{-3}$ |
| 11 | S6a,b | - | - | - | - | - | - | $5 \times 10^{-5}$ |
| 12 | S6c,d | - | - | - | - | - | - | $5 \times 10^{-5}$ |

Table 1: Vicariance parameters. Dashes match the default parameters.

| parameter settings | figure | $N_1$ | $N_2$ | $N_3$ | $m_{1\to 2}$ | $m_{2\to 1}$ | $m_{1,2\to 3}$ | $m_{3\to 1,2}$ |
|---|---|---|---|---|---|---|---|---|
| default/1 | 1, S7 | 24,300 | 2,700 | 3,000 | $5 \times 10^{-6}$ | $5 \times 10^{-6}$ | $5 \times 10^{-6}$ | $5 \times 10^{-6}$ |
| 2 | S2a,b | - | - | - | - | - | - | - |
| 3 | S2c,d | - | - | - | - | - | - | - |
| 4 | S2e,f | - | - | - | - | - | - | - |
| 5 | S3a,b | - | - | - | - | - | - | - |
| 6 | S3c,d, S8 | - | - | - | - | - | - | - |
| 7 | S3e,f | - | - | - | - | - | - | - |
| 8 | S4 | - | - | - | - | - | - | - |
| 9 | S5a,b | - | - | - | 0 | 0 | 0 | 0 |
| 10 | S5c,d | - | - | - | $5 \times 10^{-5}$ | $5 \times 10^{-5}$ | $5 \times 10^{-5}$ | $5 \times 10^{-5}$ |
| 11 | S6a,b | - | - | - | $1.85 \times 10^{-8}$ | $2.06 \times 10^{-9}$ | $1.67 \times 10^{-8}$ | $1.85 \times 10^{-9}$ |
| 12 | S6c,d | - | - | - | $1.85 \times 10^{-5}$ | $2.06 \times 10^{-6}$ | $1.67 \times 10^{-5}$ | $1.85 \times 10^{-6}$ |

Table 2: Founder event-like vicariance event parameters. Dashes indicate the parameters match the default parameters.

was less extreme ($b = 0.33$). In the "founder event-like vicariance" scenario, the migration rates match the founder event scenario and the population splits were unequal (0.9 and 0.1 of the ancestral population was allocated to the two daughter populations). The exception to this was parameter settings 4 and 5 for the migration rates, where the migration rates were chosen to keep the expected number of migrants the same (see 3.3.2 for more detail).

#### 3.1 Branch lengths and divergence times

##### 3.1.1 Branch lengths

Coalescent theory predicts that properties of gene trees, such as the probability of discordance, depend on the length of time between speciation events relative to the effective population size. Less discordance is expected with longer internal branches, as there is more time for lineages to coalesce prior to the speciation time (in backwards time), where lineages from other species allow for the possibility of discordance. However, the exact probabilities of discordance

| parameter settings | figure | $N_{s1}$ | $N_{s2}$ | $N_{s3}$ | $b$ | $m_{1\to 2}$ | $m_{2\to 1}$ | $m_{1,2\to 3}$ | $m_{3\to 1,2}$ |
|---|---|---|---|---|---|---|---|---|---|
| default/1 | 1, S7 | $10^4$ | $10^4$ | $10^4$ | $10^{-3}$ | $5 \times 10^{-6}$ | $5 \times 10^{-6}$ | $5 \times 10^{-6}$ | $5 \times 10^{-6}$ |
| 2 | S2a,b | - | - | - | - | - | - | - | - |
| 3 | S2a,b | - | - | - | - | - | - | - | - |
| 4 | S2a,b | - | - | - | - | - | - | - | - |
| 5 | S3a,b | - | - | - | - | - | - | - | - |
| 6 | S3c,d, S8 | - | - | - | - | - | - | - | - |
| 7 | S3e,f | - | - | - | - | - | - | - | - |
| 8 | S4 | $2 \times 10^4$ | $2 \times 10^4$ | $2 \times 10^4$ | - | - | - | - | - |
| 9 | S5a,b | - | - | - | - | 0 | 0 | 0 | 0 |
| 10 | S5c,d | - | - | - | - | $5 \times 10^{-5}$ | $5 \times 10^{-5}$ | $5 \times 10^{-5}$ | $5 \times 10^{-5}$ |
| 11 | S6a,b | - | - | - | - | $5 \times 10^{-9}$ | - | $5 \times 10^{-9}$ | - |
| 12 | S6c,d | - | - | - | - | - | - | - | - |

Table 3: Founder event parameters. Dashes indicate the parameters match the default parameters.

| parameter settings | figure | $N_{s1}$ | $N_{s2}$ | $N_{s3}$ | $b$ | $m_{1\to2}$ | $m_{2\to1}$ | $m_{1,2\to3}$ | $m_{3\to1,2}$ |
|---|---|---|---|---|---|---|---|---|---|
| default/1 | 1, S7 | $10^4$ | $10^4$ | $10^4$ | 0.33 | $10^{-4}$ | $10^{-4}$ | $10^{-4}$ | $10^{-4}$ |
| 2 | S2a,b | - | - | - | - | - | - | - | - |
| 3 | S2c,d | - | - | - | - | - | - | - | - |
| 4 | S2e,f | - | - | - | - | - | - | - | - |
| 5 | S3a,b | - | - | - | - | - | - | - | - |
| 6 | S3c,d, S8 | - | - | - | - | - | - | - | - |
| 7 | S3e,f | - | - | - | - | - | - | - | - |
| 8 | S4 | $2 \times 10^4$ | $2 \times 10^4$ | $2 \times 10^4$ | - | - | - | - | - |
| 9 | S5a,b | - | - | - | - | 0 | 0 | 0 | 0 |
| 10 | S5a,b | - | - | - | - | $10^{-3}$ | $10^{-3}$ | $10^{-3}$ | $10^{-3}$ |
| 11 | S6a,b | - | - | - | - | $1.5 \times 10^{-4}$ | $3 \times 10^{-4}$ | $1.5 \times 10^{-4}$ | $3 \times 10^{-4}$ |
| 12 | S6c,d | - | - | - | - | - | - | - | - |

Table 4: Vicariance event-like founder event parameters. Dashes indicate the parameters match the default parameters.

| parameter settings | figure | $\tau_{1,2}$ | $t_{1,2}$ | $\tau_{1,2,3}$ | $t_{1,2,3}$ |
|---|---|---|---|---|---|
| default/1 | 1, S7 | $2 \times 10^4$ | $3 \times 10^4$ | $4 \times 10^4$ | $5 \times 10^4$ |
| 2 | S2a,b | - | - | $6 \times 10^4$ | $7 \times 10^4$ |
| 3 | S2c,d | - | - | $10^5$ | $1.1 \times 10^5$ |
| 4 | S2e,f | - | - | $1.8 \times 10^5$ | $1.9 \times 10^5$ |
| 5 | S3a,b | $4 \times 10^4$ | $5 \times 10^4$ | $6 \times 10^4$ | $7 \times 10^4$ |
| 6 | S3c,d, S8 | $8 \times 10^4$ | $9 \times 10^4$ | $10^5$ | $1.1 \times 10^5$ |
| 7 | S3e,f | $1.6 \times 10^5$ | $1.7 \times 10^5$ | $1.8 \times 10^5$ | $1.9 \times 10^5$ |
| 8 | S4 | - | - | - | - |
| 9 | S5a,b | - | - | - | - |
| 10 | S5a,b | - | - | - | - |
| 11 | S6a,b | - | - | - | - |
| 12 | S6c,d | - | - | - | - |

Table 5: Times shared across scenarios. Dashes indicate the parameters match the default parameters.

and distribution of coalescence times are not derived for more complex models, such as the models we are using with changing population sizes and migration.

As such, the impact of the duration of $s_1s_2$ (in the species tree, Fig. S1c) on gene tree distributions was investigated with simulations. As the branch length of $s_1s_2$ increases, the proportion of discordant gene trees drops substantially for all scenarios (Fig. S2a,c,e). If branch $s_1s_2$ is long, almost all of the lineages will coalesce prior to the bottleneck (backwards in time; Fig. S2), leaving little to no information to distinguish vicariance from founder event speciation. This reflects the inherent limitations of the coalescent in teaching us about old events within one population or species.

As different scenarios were compared, we verified that the average node ages are slightly higher under the realistic founder events when the $s_1s_2$ branch is longer, as the bottleneck occurs later. For realistic vicariance, on the other hand, the average node ages are slightly younger because the population does not grow in size until later, leading to a higher coalescent rate until the population size increases. Similar to the main text, the vicariance-like founder event scenario still is intermediate the realistic scenarios. Contrasting the main text, the distributions of the founder event-like vicariance scenario are more distinct from the realistic founder event, with larger average node ages and more discordant gene trees. This happens as a result of the longer $s_1s_2$ branch, which causes population size to stay larger for longer in the founder event-like vicariance event, since $N_{1,2}$ is larger in this case. The outcome is greater coalescent times on average. The minimum and maxmimum node age distribution remain similar as the branch length of $s_1s_2$ increases (Fig. S2b,d,f).

##### 3.1.2 Divergence times

While coalescent theory predicts species tree internal branch lengths impact the probabilities of gene tree discordance and the distribution of coalescence times, the terminal branch lengths should be immaterial to discordance when each species contributes a single sample. In a simple MSC (without migration) and a single sample per species, the earliest lineages can coalesce is after the first divergence time (backward time). To confirm this intuition with our model complex models, we conducted additional simulations varying the divergence times but not the internal branch lengths.

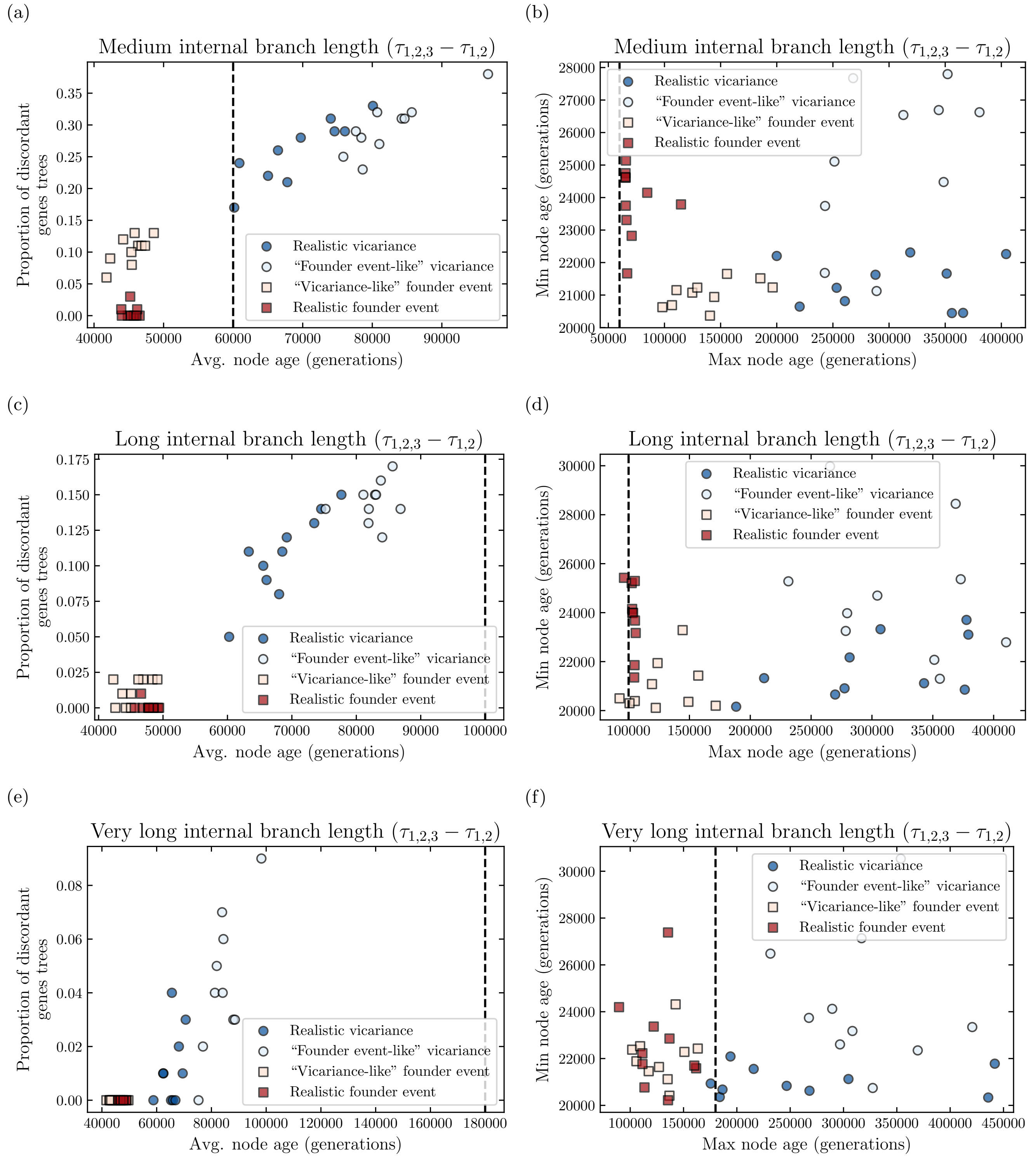


Figure S2: Dependence of coalescence times and gene tree discordance on length of branch $s_1s_2$. (a), (c), and (e) show the proportion of discordant gene trees for each replicate by the average coalescence time of the samples from $s_1$ and $s_2$. (b),(d), and (f) show the minimum and maximum coalescence times of the samples from $s_1$ and $s_2$ in each simulation. Top to bottom, internal branch length is $4 \times 10^4$, $8 \times 10^4$, and $16 \times 10^4$. In all cases, $t_{1,2} - \tau_{1,2} = 10^4$ (tables 1 to 4 parameter settings 2-4). The dashed lines show $\tau_{1,2,3}$.

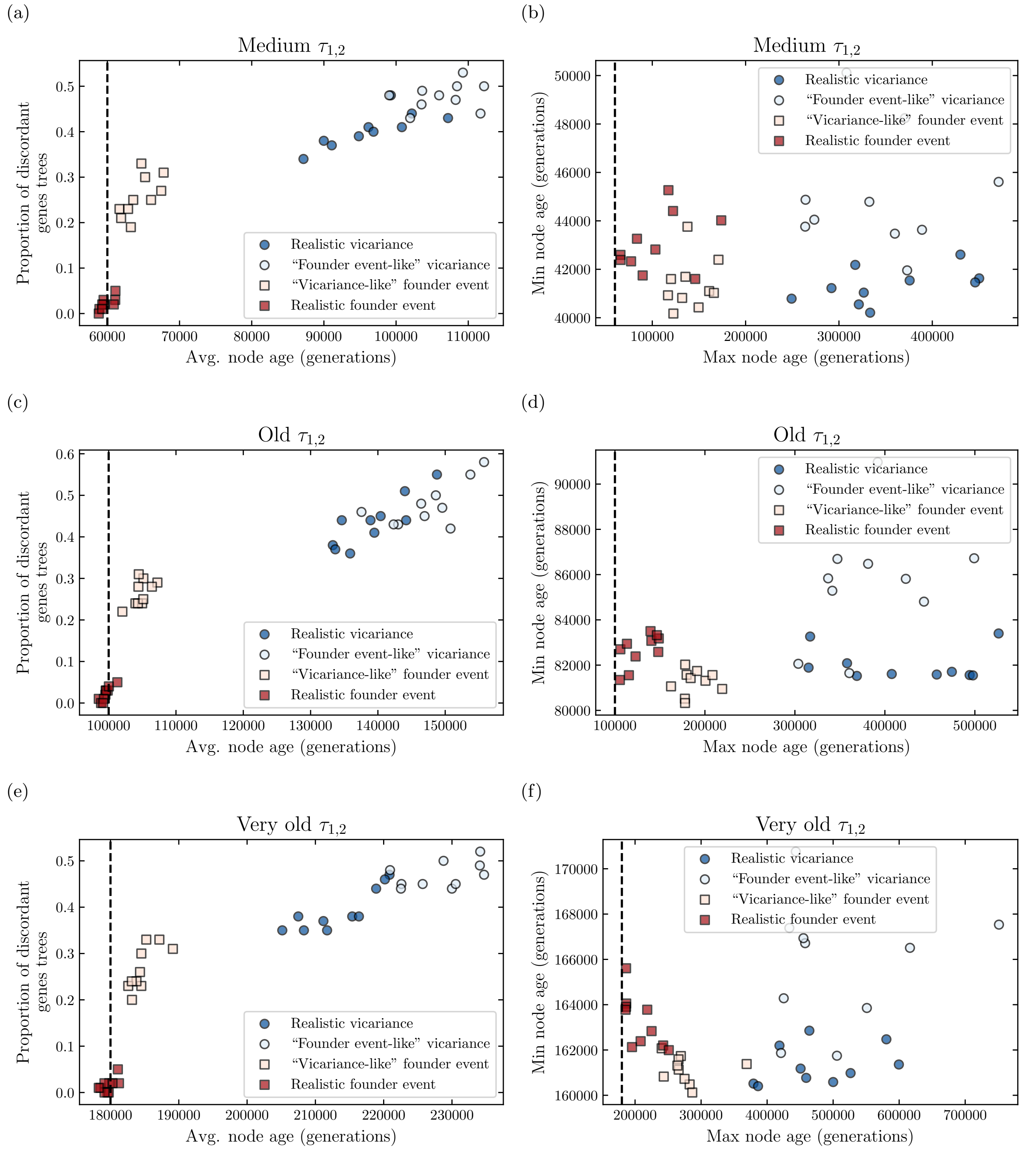


Figure S3: Dependence of coalescence times and gene tree discordance on $\tau_{1,2}$. (a), (c), and (e) show the proportion of discordant gene trees for each replicate by the average coalescence time of the samples from $s_1$ and $s_2$. (b),(d), and (f) show the minimum and maximum coalescence times of the samples from $s_1$ and $s_2$ in each simulation. Top to bottom, $\tau_{1,2}$ is $4 \times 10^4$, $8 \times 10^4$, and $16 \times 10^4$ (table 5 parameter settings 5-7). The length of branch $s_1s_2$ is constant. The dashed line show $\tau_{1,2,3}$.

As predicted, the divergence time between $s_1$ and $s_2$ ($\tau_{1,2}$) only impacts the absolute time of coalescence times, shifting the distribution by the difference in branch length of $\tau_{1,2}$, but not the coalescent times relative to the divergence time nor the frequency of gene tree discordance (Fig. S3). This is because no events can occur before two lineages are in the same population. There is no opportunity for lineages to be in the same population until $\tau_{1,2}$, when migration begins.

#### 3.2 Effective population size

Effective population sizes determine the rate of the coalescent, and thus impact both the distribution of coalescence times and the probabilities of gene tree discordance. One way that patterns could look more similar between the vicariance and founder event scenarios would be if the effective population size of $s_1s_2$ were the same, as then they would have the same coalescence rate in $s_1s_2$ (though different migration rates should impact gene tree distributions), at least until the bottleneck began in backward time.

To investigate this, simulations were conducted with matching effective population sizes for $s_1s_2$ for the realistic founder event and realistic vicariance scenarios. To make the population sizes equal across scenarios, the population size of $s_1s_2$ was increased for the founder event scenario (as opposed to decreased for the vicariance event scenario). This also increases size of $N_1$ and $N_2$ as they are all equal. Since the size of $N_1$, $N_2$, and $N_{1,2}$ are larger in comparison to the main text for the founder event scenarios, the distributions of coalescent times shift to be larger (Fig. S4 in comparison to Fig. 2). The bottleneck size is a proportion of the original population size, so the population size at the bottleneck is larger in comparison to the main text, allowing more lineages to escape the bottleneck. Additionally, the population size is larger longer with a larger initial population size (comparing parameter settings 1 and 12). This means that the bottleneck causes coalescent events slightly later when the population size is larger (Fig. S4a in comparison to Fig. 2a). Despite some quantitative shifts, the general patterns remain qualitatively similar between the scenarios. This suggests that difference in gene tree distributions may persist between founder event and vicariance scenarios even if demographics of the ancestral populations are more similar.

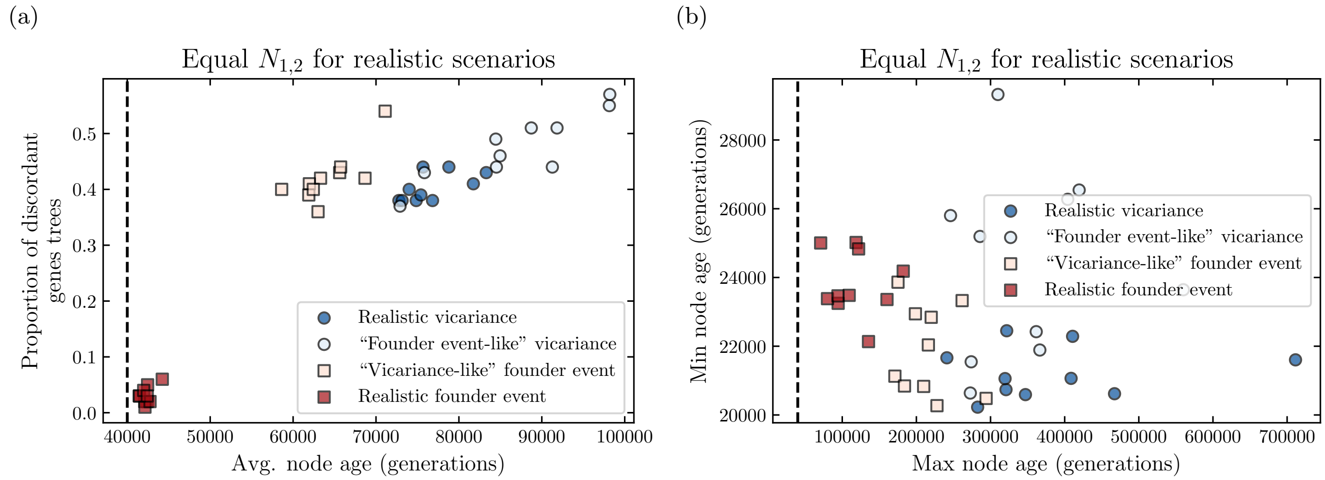


Figure S4: Impact of population size of $s_1s_2$ on coalescence times and gene tree discordance. (a) shows the proportion of discordant gene trees for each replicate by the average coalescence time of the samples from $s_1$ and $s_2$. (b) shows the minimum and maximum coalescence times of the samples from $s_1$ and $s_2$ in each simulation. The populations sizes in $s_1s_2$ are the same for the realistic founder and realistic vicariance (tables 1 and 3 parameter setting 8). The dashed lines show $\tau_{1,2,3} = 4 \times 10^4$.

#### 3.3 Migration

##### 3.3.1 Migration rates

Migration has a large impact on gene tree distributions. With high enough migration, populations from the same species are effectively panmictic, meaning coalescence events should be equally likely between samples from the same or different populations. On the other end of the spectrum, no within-species migration after population divergence means the earliest coalescent times must happen, forward in time, before *populations* begin to diverge, as opposed to just before the speciation completion time. Empirically, there is likely a broad range of how much migration occurs during the process of speciation.

As such, the impact of no migration and higher migration rates on gene tree distributions was investigated with simulations. There are two main impacts of migration in our simulations. First, it allows lineages to coalesce earlier (backward time), during the period of population subdivision by migrating to a population with another lineage. Otherwise without migration, there is only one lineage in each population until $t_{1,2}$. Second, if a lineage migrates to a different population without a bottleneck, it may escape the effect of a bottleneck in the original population in the founder event scenarios. With that in mind, consider first the case of no migration. Without migration, the populations are completely isolated as soon as there is population subdivision so the population divergence times are effectively older, occurring at $t_{1,2,3}$ instead of $\tau_{1,2,3}$ and $t_{1,2}$ instead of $\tau_{1,2}$. Since migration allows for lineages to coalesce during the period with population subdivision for the simulations in the main text, without migration the minimum gene tree node ages are older in comparison for all scenarios (Fig. S5b comparing to Fig. 2b). However, the coalescence times are not always older (Fig. S5a,b comparing to Fig. 2). Without migration, both lineages from $\mathrm{s}_1$ and $\mathrm{s}_2$ must go through the bottleneck in population $\mathrm{s}_1\mathrm{s}_2$, which forces the lineages to coalesce as the population shrinks (Fig. S5b, maximum node age), leading to the maximum nodes ages being smaller without migration in many cases. Despite changes in the exact times of average, minimum, and maximum coalescence events and slight changes in the proportions of discordant gene trees, the general patterns and ability to distinguish the distributions of the scenarios remain similar.

With high migration rates, the average node age is somewhat lower for the realistic founder scenario and the proportion of discordant gene trees is slightly higher (Fig. S5c compared to Fig. 2a and S5a). Higher migration causes a greater proportion of lineages to move between populations $\mathrm{s}_1$ and $\mathrm{s}_2$, and thus coalesce during the period with population structure. However, the maximum node age is higher with high migration (Fig. S5d in comparison to Fig. 2b and S5b), since migration allows the lineages to move to $\mathrm{s}_3$, where there is no bottleneck. The realistic founder has lower migration rates than the "vicariance like" founder event, explaining the lower minimum coalescent times for the latter. In summary, while migration rates have a large impact on the gene tree and coalescent time distributions, the distributions for vicariance and found events are still distinct even with rather extreme migration rates.

###### 3.3.2 Migration in expected number of migrants

In the original simulation, the migration rates were symmetric with respect to the proportion of the destination population replaced (forward in time). This can lead to really extreme values, particularly if effective population size is changing. For example, $\mathrm{s}_2$ has a large proportion (in the population genetics sense, though possibly still a small number) of its population replaced through migration each generation but $\mathrm{s}_1$ is small, this may require a very large proportion of $\mathrm{s}_1$ to migrate. An alternative to symmetric migration in term of the proportion replaced would be to use migration that is symmetric in term of the expected number of migrants.

To investigate whether the simulation results were sensitive to the migration parameterized, the migration rates were varied such that the rates were symmetric in terms of the expected number of migrants between the populations exchanging migrants. This is possible for the vicariance scenarios, where the population sizes are constant within a population. However, since the MSPRIME simulator is parametrized in terms of the proportion replaced, the expected number of migrants changes overtime as the population size changes. Thus, cases were examined where the expected number of migrants were symmetric between populations before or after the bottleneck. To determine the expected number of migrants from population $i$ to $j$ in forward time, $M_{i,j}$, $M_{i,j} = N_j m_{i,j}$, where $m_{i,j}$ is the proportion of population $j$ replaced by population $i$ each generation forward in time.

When migration is symmetric in expected number of migrants before or after the bottleneck, the results look similar to when the expected proportion of the descendant population replaced is symmetric (Fig. S6 comparing to Fig. 2). When they are symmetric in expected number of migrants before the bottleneck only, the "founder event-like" vicariance event has less variance in the minimum age and a smaller minimum age on average (Fig. S6d comparing to Fig. 2b). This is due to the much larger migration rate. Note that some of the migration rates are likely unrealistically high and some of them are very low (tables 1 to 4 scenarios 11 and 12). These results suggest that the general patterns in the main text are not sensitive to how migration is parametrized, even with extreme rates.

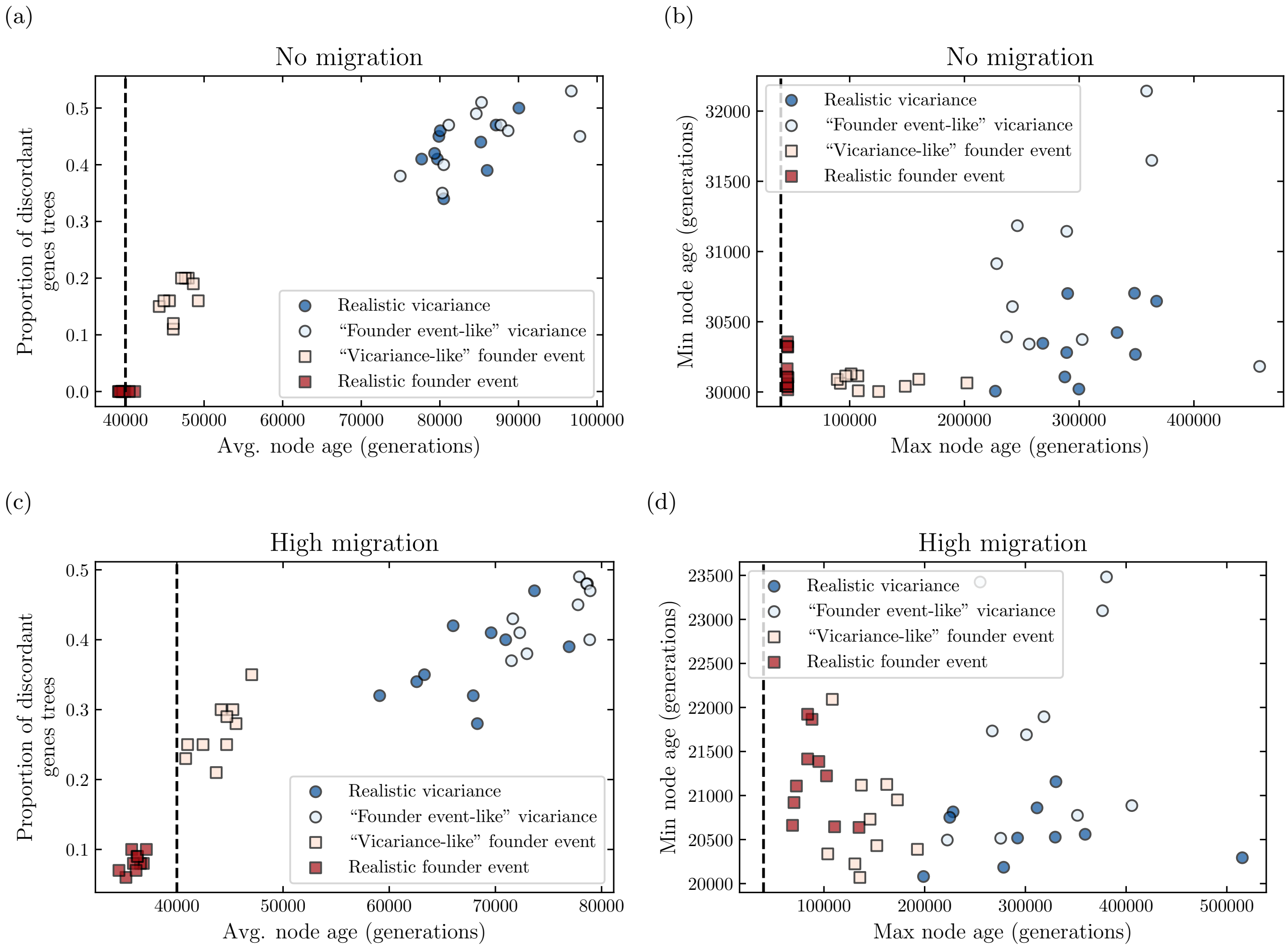


Figure S5: Dependence of coalescence times and gene tree discordance on migration rates. (a) and (c) show the proportion of discordant gene trees for each replicate by the average coalescence time of the samples from $s_1$ and $s_2$. (b) and (d) show the minimum and maximum coalescence times of the samples from $s_1$ and $s_2$ in each simulation. (a) and (b) have no migration. (c) and (d) have high migration rates (tables 1 to 4 parameter settings 9 and 10). These rates are an order of magnitude higher than in Fig. 2 in the main text. The dashed line show $\tau_{1,2,3} = 4 \times 10^4$, which is when migration begins between $s_1 s_2$ and $s_3$ (if it is non-zero) and when the population size begins decreasing (backward in time) for the bottleneck in $s_2$.

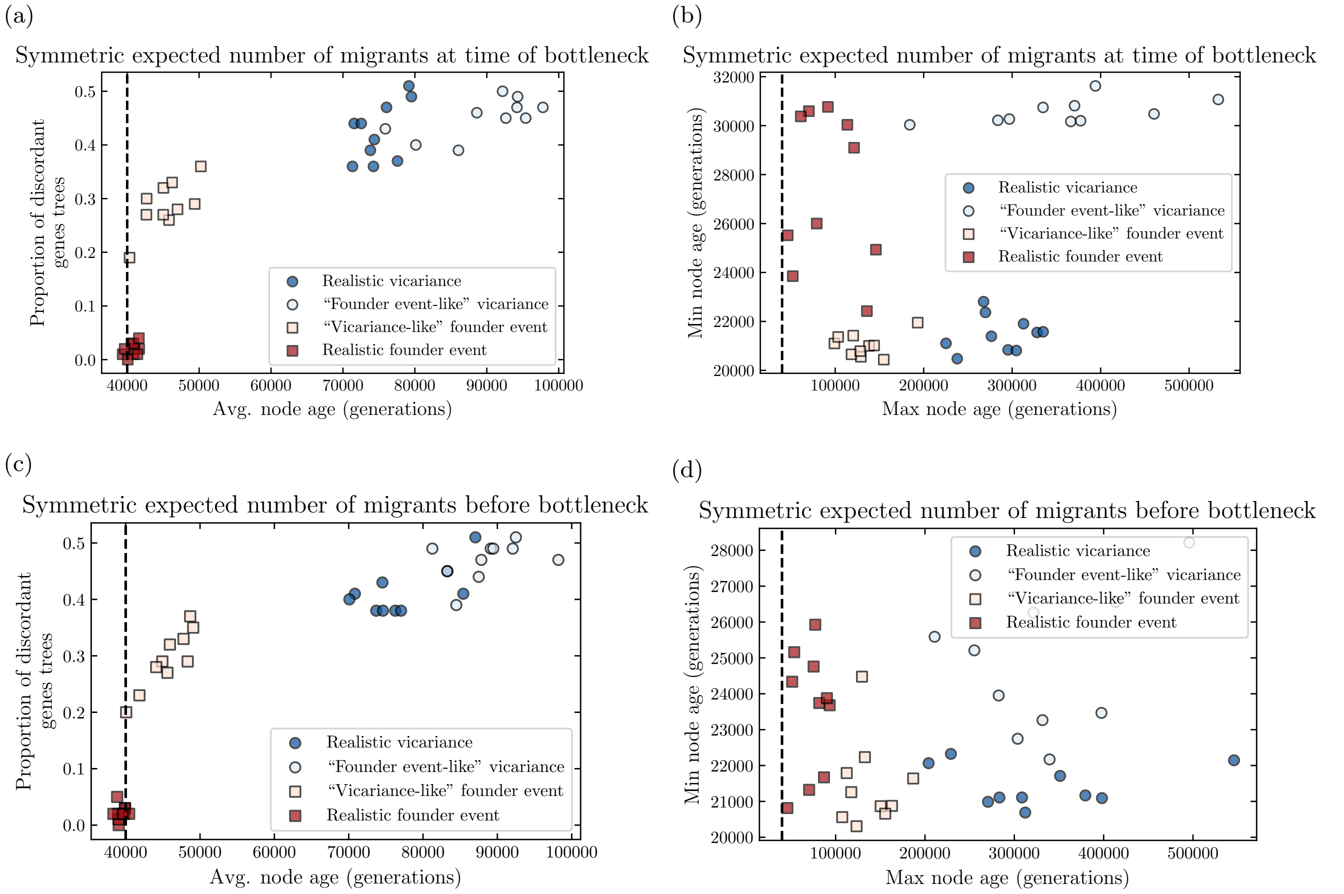


Figure S6: Sensitivity of distributions of coalescence times and gene tree discordance to alternative parametrization of migration. The expected number of migrants is equal to 1.0 migrants per generation and is the same in each direction for the realistic vicariance scenario in all panels. For the realistic founder event, (a) and (b) the expected number of migrants is the same in each direction at the time of the bottleneck and it equal to $5 \times 10^{-5}$ migrants per generation. For the realistic founder event in (c) and (d), the expected number of migrants in each direction is the same before the bottleneck (backward time) and it equal to $5 \times 10^{-2}$ migrants per generation. Since the proportion replaced is constant, this means that there will be more migrants into the new population going forward in time (tables 1 to 4 parameter settings 11 and 12).

#### 3.4 Impact of sample size per species

The main text results suggest gene trees distributions could be used to study deeper speciation events in a phylogeny, but these simulations provide only limited information about the speciation event that gave rise to the two extant sister species. This is due to the fact that multiple samples per species are necessary to see the impact of demography within a species, i.e. there is no signature different demographic histories such as the bottleneck in $s_1$ with a single sample in $s_1$.

To investigate the history of divergence between $s_1$ and $s_2$, consider sampling two lineages from $s_1$ and one lineage from $s_2$ for 100 loci. With two samples from $s_1$, there will be coalescent events within the species $s_1$, thereby informing its demographic history. Now we consider the coalescent times between the two samples from $s_1$, rather than the coalescent times between the samples from $s_1$ and $s_2$ as was previously considered. As before, the bottleneck in the founder event scenario forces the lineages, this time from $s_1$, to coalesce before (backwards in time) reaching the ancestral population, leading to less discordance and smaller coalescent times in that scenario relative to the vicariance scenario (Fig. S7a). Conversely, consider repeating the simulation with one sample from $s_1$ and two samples from $s_2$, and comparing the coalescent times for the lineages from $s_2$. The major difference between this and sampling two lineages from $s_1$ the absence of a bottleneck in $s_2$. The realistic vicariance coalescence times and amount of gene tree discordance remains similar (comparing Fig. S7a and b), but the realistic founder event coalescence times are much older with much more discordant gene trees resulting from the lack of a bottleneck. The coalescence times are still smaller on average in the realistic founder event in comparison to the realistic vicariance because the population sizes for $s_1s_2$ differ. The proportion of discordant genes trees is also smaller for the realistic founder event. Since the migration rate is lower, lineages are more likely to stay in the same population and coalesce during the period with substructure.

These results suggest that there should be information to distinguish different types of speciation events that give rise to two extant sister species in some cases. Additionally, in the case of a founder event, it may be possible to distinguish which of the two populations was the original population and which arose as the result of a founder event.

##### 3.4.1 Branch lengths and divergence times

While the divergence time of $s_1$ and $s_2$ does not impact the relative distribution of coalescence times or distribution of gene tree discordance with a single sample from each species, this may not be the case with multiple samples from either species $s_1$ or $s_2$ since lineages may coalesce before $\tau_{1,2}$.

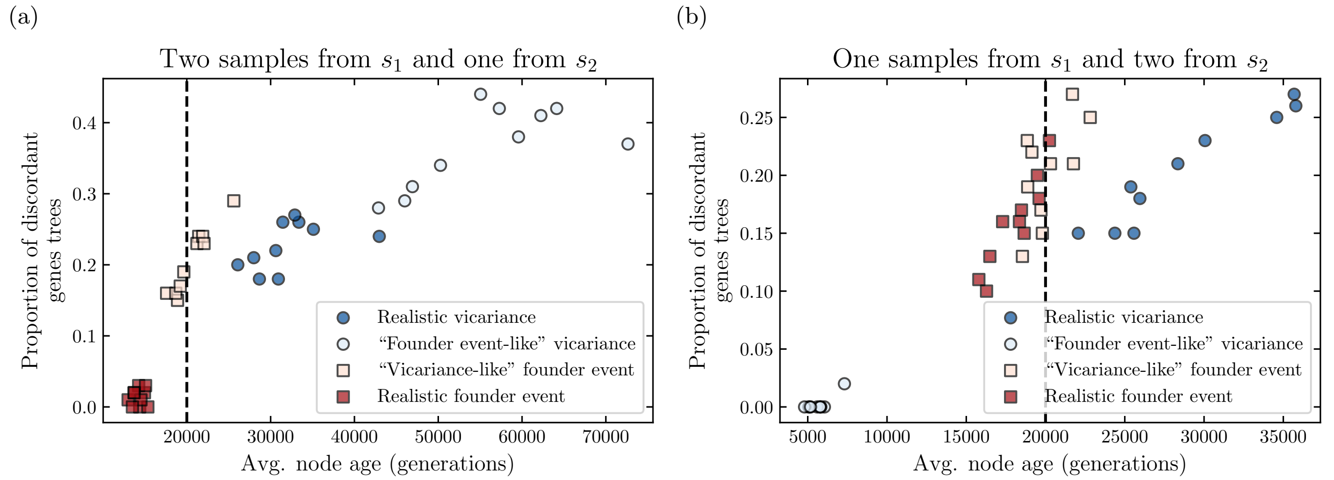


Figure S7: Distributions of coalescent times and discordant gene trees with multiple samples from the same population. The x-axis shows the average coalescent time of two samples from $s_1$ and the average coalescent time of two samples from $s_2$ for (a) and (b), respectively. The y-axis shows the proportion of gene trees where the samples from the same daughter population are not monophyletic. (a) has two samples from $s_1$, one sample from $s_2$, and no samples from $s_3$. (b) has one sample from $s_1$, two samples from $s_2$, and no samples from $s_3$. Each simulation has 100 gene trees. There are 10 replicates of each scenario. The parameters are the same as in Fig. 1. The dashed lines show $\tau_{1,2}$.

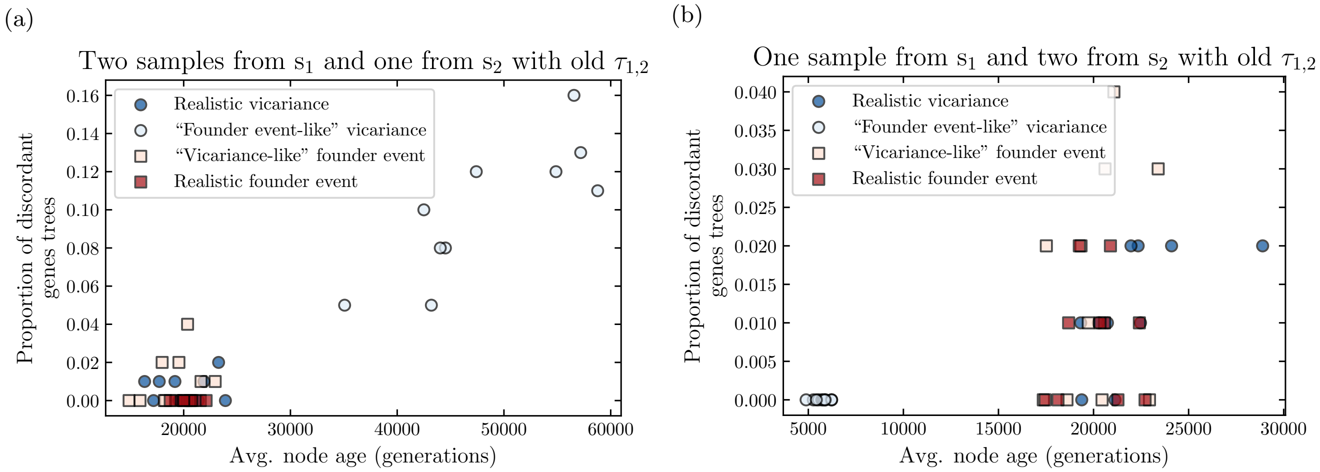


Figure S8: Impact of branch $\tau_{1,2}$ on distribution of coalescence times and gene tree discordance with multiple samples from $s_1$ or $s_2$. (a) shows the proportion of gene trees where the samples $s_1$ are not monophyletic by the average coalescence time of the two samples from $s_1$ for simulations with two samples from $s_1$ and one sample from $s_2$. (b) shows the proportion of gene trees where the samples $s_2$ are not monophyletic by the average coalescence time of the one sample from $s_1$ for simulations with two samples from $s_2$ and two samples from $s_2$. $\tau_{1,2}$ is $8 \times 10^4$ (table 5 parameter setting 10).

The time of population divergence $s_1$ and $s_2$, $\tau_{1,2}$, does impact the distribution of coalescent times when there are multiple samples from $s_1$ or $s_2$ (Fig. S8). As $\tau_{1,2}$ increases, the proportion of trees where the samples from the same species are not monophyletic drops substantially, since there is more time for samples within a species to coalesce before any lineages from a different species are in the same population. The coalescent times for the vicariance scenarios also decrease on average, as the population size remains smaller longer, so the coalescent rate remains high for longer. This suggests that with sister taxa, older divergence times limit the ability to distinguish different types of speciation scenarios.
